# Temperature dependent response of microcystin-LR in acclimated *Microcystis aeruginosa:* highest content expected near the growth optimum

**DOI:** 10.1101/2025.06.20.660678

**Authors:** Pierre-Louis Lalloué, Clarisse Mallet, Alexandre Bec, Apostolos-Manuel Koussoroplis, Fanny Perrière, Delphine Latour

**Affiliations:** Université Clermont Auvergne, CNRS, Laboratoire Microorganismes: Génome Environnement, Clermont–Ferrand, France

**Keywords:** Cyanobacteria, *Microcystis aeruginosa*, microcystin, temperature, acclimated cells, thermal performance curves, cell volume

## Abstract

As climate change raises global temperatures and increases the frequency of cyanobacterial blooms, understanding how rising mean temperatures affect cyanotoxin content is crucial. However, no clear consensus exists, and the use of different methodologies, including different units of measurement and experimental conditions could significantly alter the yield of the relationship between temperature and toxins content. In this study, we assessed free microcystin content and cell volume in *Microcystis aeruginosa* PCC 7806 acclimated to seven temperatures spanning its entire thermal niche. This experimental design firstly highlighted the significant reduction in cell volume with rising temperatures between 17°C and 29°C. As a result, when the microcystin concentration was normalized by its cell volume, its temperature response was transformed from a negative correlation to a bell-shaped curve, with higher free MC-LR content measured at an estimated optimum temperature of 26.2°C, close to the thermal growth optimum of *Microcystis aeruginosa*. These findings provide new insights into the effects of climate warming on microcystin content.

**Importance:** Microcystin-LR is a widespread cyanotoxin, originally known for its liver toxicity. In freshwater environments, cyanotoxins are an increasing concern as harmful cyanobacterial blooms become more frequent with rising global temperatures. *Microcystis aeruginosa*, a common bloom-forming species found worldwide, is a major producer of microcystin-LR. Understanding how environmental factors such as temperature influence toxin content in this species is essential for predicting bloom toxicity under future climate scenarios. However, current knowledge remains fragmented due to numerous factors that can influence its production and also to different way of measuring toxins and expressing their concentrations (cell or µm3). Confirming that temperature greatly modifies biovolume of *M. aeruginosa*, this study offers new insights by highlighting the importance of considering cell volume when evaluating toxin content. Integrating changes in cell size helps reconcile earlier conflicting results and contributes to a more accurate understanding of how temperature affects toxin production in cyanobacteria.

**Graphical abstract:** 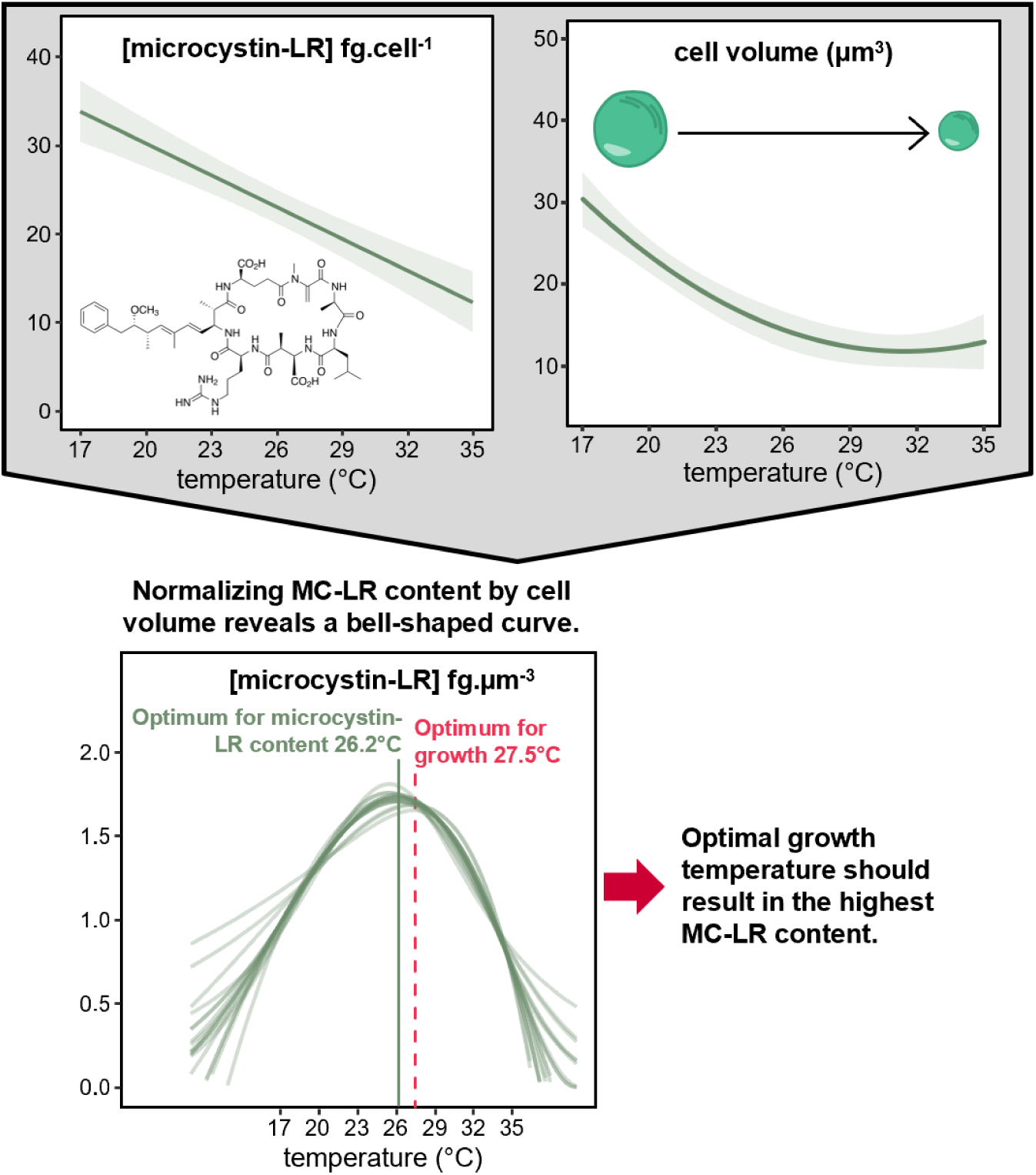

## 1. Introduction

Cyanobacterial blooms pose a global threat to freshwater ecosystems, causing adverse impacts on aquatic trophic networks, the economy, and human health (Huisman et al., 2018). A key concern associated with these blooms is the production of cyanotoxins, which can affect organisms at all trophic levels from eukaryotic primary producers (Teneva et al., 2023) and zooplanktonic primary consumers (Freitas et al., 2014) to predatory fish (Shahmohamadloo et al., 2021). Beyond their ecological impacts, cyanotoxins can directly harm human health if contamination of recreational or drinking water reservoirs occurs. Moreover, their presence imposes significant economic burdens, including costs related to public health, fisheries, tourism, recreational activities, and water quality monitoring and management (Carmichael and Boyer 2016). Among the cyanotoxins, microcystins are the most studied and ubiquitous. Primarily known as hepatotoxins inhibiting the protein phosphatase 1 and 2A in hepatocytes, they seem to have a larger spectrum. Some studies have found the effects of microcystins to include being spermatotoxic, cancerogenic and linked to enhancing Alzheimer’s, causing a growing concern for human health (Chen et al., 2011; Li et al., 2012; Lone et al., 2015).

The frequency of harmful cyanobacterial blooms has increased during the Anthropocene, driven in part by the eutrophication of aquatic ecosystems and exacerbated by global warming (Huisman et al., 2018). Cyanobacteria, which typically have higher thermal optima than other phytoplankton groups and benefit from prolonged lake stratification, are expected to bloom more frequently as surface temperatures continue to rise (Huisman et al., 2018). However, the effect of temperature on the microcystin content of cyanobacterial cells remains puzzling, making it difficult to predict the toxicity of future blooms in a warming world. Despite many previous works having addressed the effects of temperature on microcystin content, there is currently no clear pattern. Indeed, while older studies report that the relationship between *Microcystis* toxin content and temperature may follow a bell-shaped curve (Sivonen et al., 1990; Van der Westhuizen & Eloff, 1985), more recent research presents contradictory findings, suggesting a linear decrease in microcystin content as temperature rises (Bui et al., 2018; Martin et al., 2020; Melina Celeste et al., 2017; Peng et al., 2018). The discrepancy between these studies could arise from culture conditions, cyanobacterial strains but also the units used to express toxin content. While recent studies typically normalize microcystin-LR (MC-LR) content by cell density, older ones often use dry weight. This distinction is critical, as cyanobacterial cell volume—and consequently dry weight—can vary with temperature (Martin et al., 2020). This difference in units could significantly influence the observed trends and their subsequent interpretations; neglecting to account for it might lead to erroneous predictions of toxin production and misrepresent the physiological role of microcystins.

To improve predictions regarding the impacts of climate change-induced temperature increases it is crucial to first establish the reaction norm of microcystin content under stable conditions. This is why, in this study, we chose to investigate the effects of temperature on the MC-LR content of *Microcystis aeruginosa* PCC7806 — a well-known and extensively studied strain recognized for its ability to produce MC-LR. To this end, the strain was acclimated to seven temperatures ranging from 17°C to 35°C, covering its full thermal niche under our culture conditions. (Acclimation at 14°C and 38°C was not achievable, as cultures collapsed at these extremes.). For the first time, our approach combines acclimation with cell volume measurements across the full thermal niche, based on the hypothesis that temperature-induced changes in cell volume could significantly influence observed trends in microcystin content. In parallel, growth rate is another temperature-dependent parameter that could, in turn, influence MC-LR content. A positive correlation between microcystin content and growth rate has already been demonstrated under nitrogen limitation (Long et al., 2001), likely due to the high nitrogen demand for MC-LR production. However, data on the effects of temperature on MC-LR content remain limited, and this relationship may reflect a trade-off between the energetic cost of MC-LR synthesis and its potential benefits for the cell. To investigate this relationship, we employed thermal performance curves (TPC) to model growth rate responses to temperature, allowing us to determine the growth optimum.

## 2. Materials and methods

### 2.1 Strains and culture condition

Cultures of *Microcystis aeruginosa* PCC 7806 were obtained from the Pasteur Culture Collection (PCC) and cultivated in WC growth media (Guillard and Lorenzen, 1972). The cultures were grown in 500 ml culture flasks, each containing 200 ml of the medium. Continuous light at 25 µmol photons.m^-2^.s^-1^ was supplied by full spectrum 15-W LEDs (380-750 nm, Aquatlantis, Portugal). Before the experiments, cultures were maintained at 20°C in temperature-controlled incubators (Pol-eko, Poland) with 90 rpm using a Stuart Orbital Shaker SSL1 (Stuart Equipment, UK).

### 2.2 Experimental design

All conditions were tested in quadruplicate batch cultures. Seven temperatures (17, 20, 23, 26, 29, 32 and 35°C) were tested, and temperatures were monitored every 5 seconds using a Hobo Pendant MX TempLogger® (OnSet Computer Corporation, Massachusetts, USA) submerged in the same volume of water as the cultures. Prior to each experiment, cultures were acclimated to the respective temperature for a period of at least two weeks (Fey et al., 2021). During this acclimation period, the cultures were maintained in exponential growth phase by diluting them twice a week with fresh WC medium. On day 0 (inoculation), cultures were diluted to a cell density of 100,000 cells.ml^-1^. To ensure that nutrient limitation did not occur, cultures were harvested for MC-LR quantification and cell volumes measurements during the mid-exponential phase, approximately two days after the onset of exponential growth (Figure S1). Sampling was performed every two days to monitor cell density.

### 2.3 Cell density and growth rate estimations

Cell density was monitored throughout the experiment by measuring optical density at 680 nm every two days using a spectrophotometer (Multiscan SkyHigh, Thermo Scientific, Massachusetts, USA). The optical density values were then converted to cell density using calibration curves. Exponential growth rates were estimated as the slope of the relationship between the log of cell density and time using the function “get.growth.rate” in the R package “growthtools” with the “sat” method (which assumes the presence of a saturation phase).

### 2.4 Intracellular MC-LR

#### 2.4.1 Intracellular MC-LR extraction

Five ml of each culture was collected, centrifuged, and stored at -80°C before freeze-drying. Toxins were then extracted from the cell pellet using 2 ml High Performance Liquid Chromatography (HPLC) grade methanol 75%, followed by ultrasonication (120 Sonic Dismembrator, Fisher Scientific, Massachusetts, USA; 15seconds/3 cycles/50% power). The extracts were centrifuged at 11,000 x g for 10 minutes at 4°C, and the supernatant was transferred into a glass test tube. Each cell pellet underwent this protocol three times and the resulting supernatants were pooled and evaporated under a nitrogen flow. The dried extracts were stored at 4°C, then resuspended in 500 µl of 20% HPLC grade methanol, and transferred into an autosampler vial for HPLC analysis.

#### 2.4.2 HPLC analysis

Microcystin-LR was quantified using the high-performance liquid chromatography with photodiode array detection (HPLC-DAD, 1260 Infinity II system, Agilent, California, USA), equipped with a Kinetex C18 column (150 x 4.6 mm, 5 µm pore size, Phenomenex Inc, California, USA). The mobile phase consisted of ultrapure water with 0.05% TFA (Buffer A) and HPLC grade acetonitrile with 0.05% TFA (Buffer B), following the gradient: starting with 20% B; 0-10 min linear increase in B from 20 to 30%; 10-25 min linear increase to 70% B; 25-26 min linear increase to 100% B; 26-30 min maintained at 100% B; 30-35 min linear decrease to 20% B. B was then maintained at 20% for 5 min allowing the column to equilibrate before the next sample. The column temperature was set at 30°C, flow rate was 1 ml.min^-1^ and the injected volume was 50 µl. To determine MC-LR concentration, the peak area at 238 nm was measured and subsequently converted into concentration using a calibration curve established from a dilution series of an external MC-LR standard (Eurofins Abraxis, Pennsylvania, USA).

### 2.5 Cell volume estimation

The volume of *M. aeruginosa* cells was estimated from cells harvested simultaneously for MC-LR analysis. These cells were fixed with 2% Lugol’s solution and stored at 4°C until analysis which was performed within the following two weeks. For each experimental condition, ∼100 cells were measured across four replicates. The measurements were performed by ZEN 3.0 software (Zeiss, Germany) on micrographs taken at 40x magnification. We assumed that cells were not perfectly spherical; therefore, we measured both a short diameter (d) and a long axis (h) for each cell. The cell volume (v) was then calculated using the following formula: 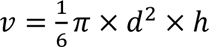 (Napiórkowska-Krzebietke and Kobos, 2016)

### 2.6 Statistical analysis

MC-LR concentration data were normalized to either cell concentration or total biovolume (cell concentration * mean cell volume) before statistical analysis. To assess the effect of temperature on growth rates and microcystin concentrations, a one-way ANOVA followed by Tukey’s post hoc test was conducted. A Pearson correlation test was used to evaluate the relationship between exponential growth rate and MC-LR concentration per volume unit. To examine the effect of temperature on cell volume, a Kruskal-Wallis test followed by Dunn’s post hoc test was performed. The assumptions of normality and homoscedasticity were verified prior to statistical analysis using the Shapiro-Wilk test for normality and Bartlett’s test for homoscedasticity, respectively. A significance threshold of p = 0.05 was applied to all statistical tests. Statistical analyses were conducted using R software (v4.1.2; R Core Team 2021).

TPC for exponential growth rates and volumetric concentrations were fitted using the rTPC package in combination with nls_multstart to estimate the initial parameters of the models (Padfield et al., 2021). We attempted to fit all the models available in the rTPC package and retained those with an R² value above 0.5 and whose initial parameter estimations successfully converged (Kontopoulos et al., 2023).

## 3. Results

### 3.1 Exponential growth rates

There were significant differences in the exponential growth rates of *M. aeruginosa* (Figure 1) depending on the test temperature, with cultures at 17°C showing the lowest rates (0.16 ± 0.02 d⁻¹), while those at 26°C and 29°C (0.52 ± 0.04 and 0.50 ± 0.05 d⁻¹, respectively) exhibited the highest rates. It should be noted that there are no significant differences between the treatments at 20, 23, 32 and 35°C. Thus, the relationship between temperature and growth rate for *M. aeruginosa* appears to follow a classical thermal performance bell-shaped curve. In this way, the 15 TPC models fitted to the growth rates data provide a good prediction of how temperature affects the growth of *M. aeruginosa,* with a mean R^2^=0.80 (Table S1). This allows us to estimate the optimal temperature (Topt) for growth at 27.5 ± 0.8°C.

**Figure 1.**
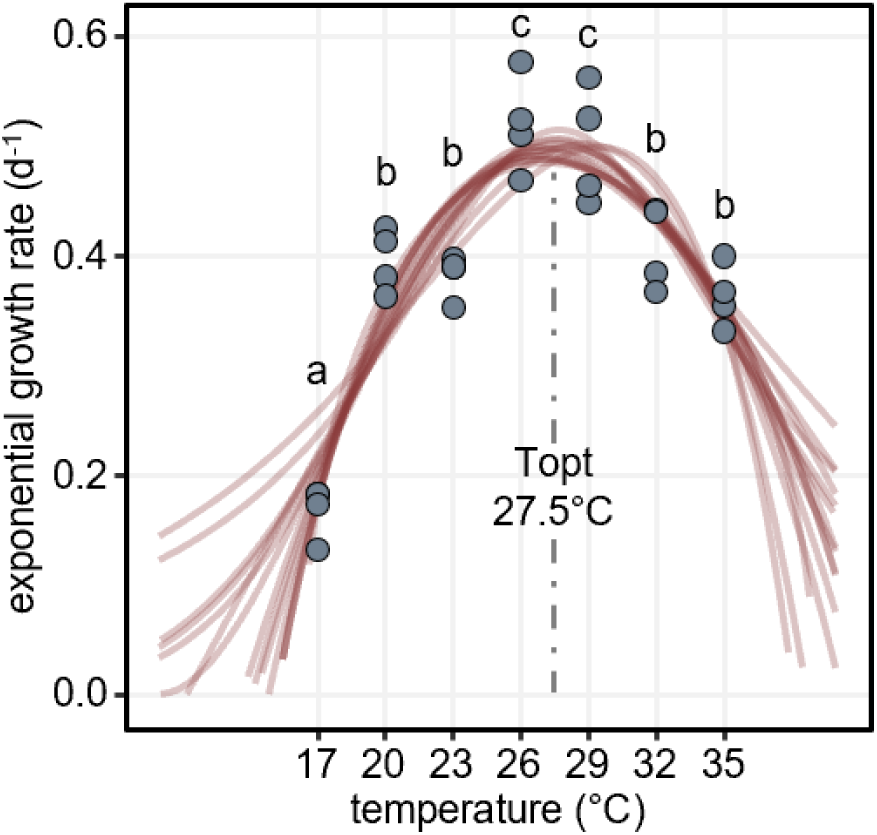
Exponential growth rates of M. aeruginosa PCC7806 as a function of temperature (n=4). The red lines depict the 15 TPC models fitted using rTPC, while the vertical grey dashed line represents the mean optimal temperature estimated from these models. Different letters indicate significant differences between temperature conditions (Table S2; ANOVA, p=1.37e-10).

### 3.2 Cell volumes

*M. aeruginosa* cell volume decreases significantly with increasing temperature, with values at each temperature being significantly lower than those at the preceding temperature between 17 and 29°C (Figure 2a). Specifically, we observed a 2.5-fold reduction in cell volume from 17°C (31.7 ± 6.6 µm^3^) to 29°C (12.5 ± 2.1 µm^3^). No significant size differences were observed between conditions at temperatures higher than 29°C. Due to this asymptote, a quadratic regression fits our data better than a simple linear regression or even a logarithmic model. The size distribution is wider at 17°C, then decreases rapidly and becomes narrower at higher temperatures (figure 2b).

**Figure 2.**
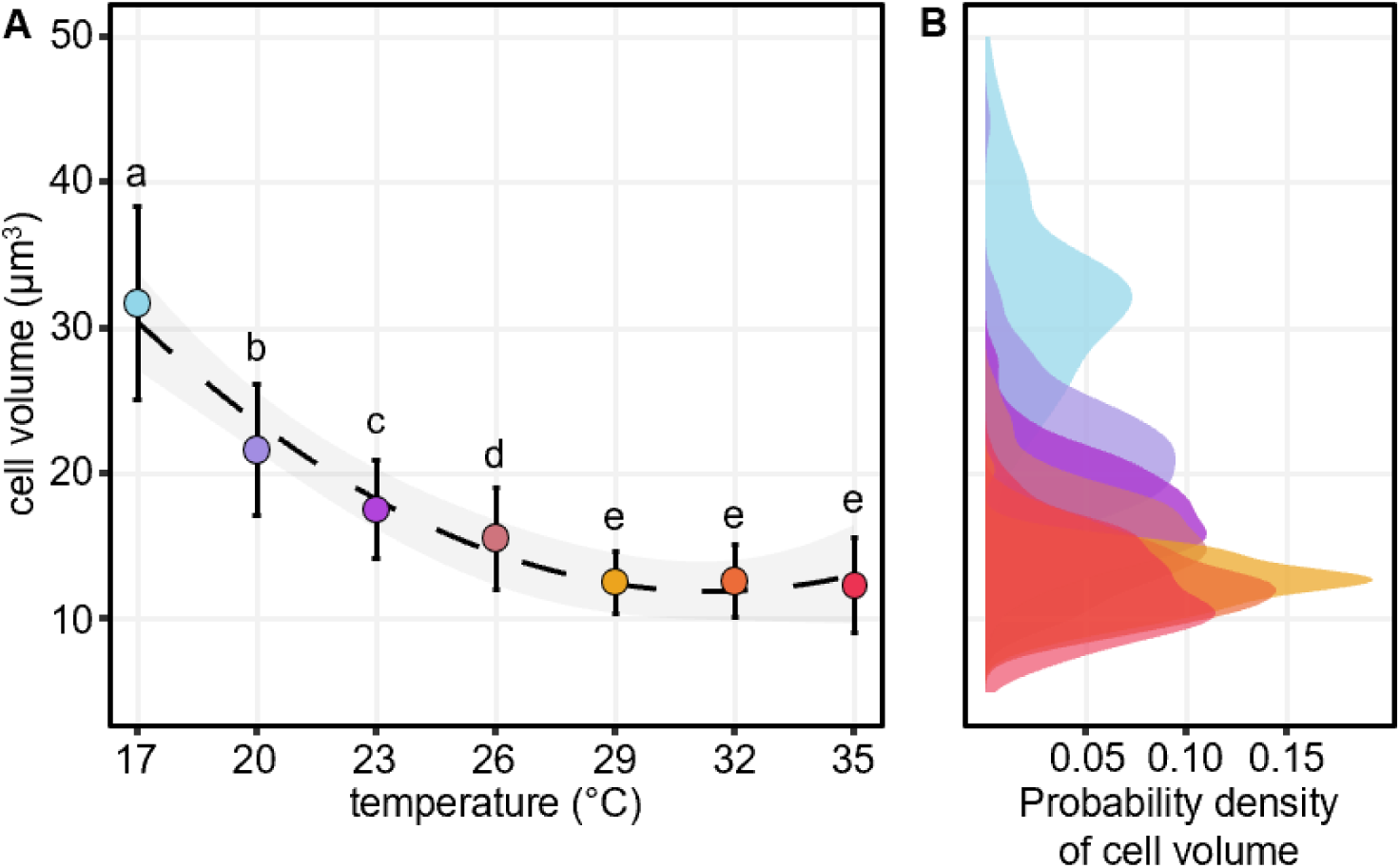
Cell volumes of M. aeruginosa PCC7806 grown at seven temperatures. **(a)** Mean cell volumes (circles) versus temperature (n∼100) with error bars set as standard deviation. The dashed line is the quadratic regression of cell volume as a function of temperature (R2=0.698, p<2e-16). Different letters indicate significant differences between temperature conditions (Table S3; Kruskal-Wallis, p<2e-16). **(b)** Distribution density of Microcystis cell volumes in the tested temperature conditions. Colors are the same as points in Figure 2. a and corresponded to the experiment temperature.

### 3.3 Temperature effects on microcystin-LR content

The intracellular concentrations of MC-LR normalized by cell density at each tested temperature are shown in Figure 3a. The concentration decreases approximately 4-fold, from 32.7 ± 3.9 fg.cell^-1^ at 17°C to 8.2 ± 0.4 fg.cell^-1^ at 35°C. There is a strong linear relationship linking cellular MC-LR concentration to temperature, as indicated by the fitted linear model (R^2^=0.7, p=3.6e-8). Between 17°C and 26°C, the only significant difference is between 17°C and 23°C (p=0.047). However, since cell volume decreases significantly with rising temperatures, the intracellular concentration of MC-LR was normalized by the mean cell volume (Figure 3b). Expressed this way, the pattern of these results shifts from a linear model to a TPC shape. The values of MC-LR content increase from 17°C to a peak at 26°C (1.9 ± 0.1 fg.µm^-3^) and then decrease to a minimum at 35°C (0.6 ± 0.03 fg.µm^-3^). A total of 19 TPC models were successfully fitted to these values, obtaining a mean R^2^=0.67 and an estimation of the optimal temperature of 26.2 ± 0.6°C for the higher production of MC-LR (Table S4). This value is slightly lower than the estimated optimal temperature for growth. The relationship between MC-LR concentration per volume and exponential growth rate is shown in Figure 3c. When considering the full temperature range, no significant correlation is observed (p = 0.166). However, this lack of significance is primarily due to the data points at 35°C. Between 17°C and 32°C, a significant positive correlation emerges between MC-LR concentration and growth rate (p = 0.012).

**Figure 3.**
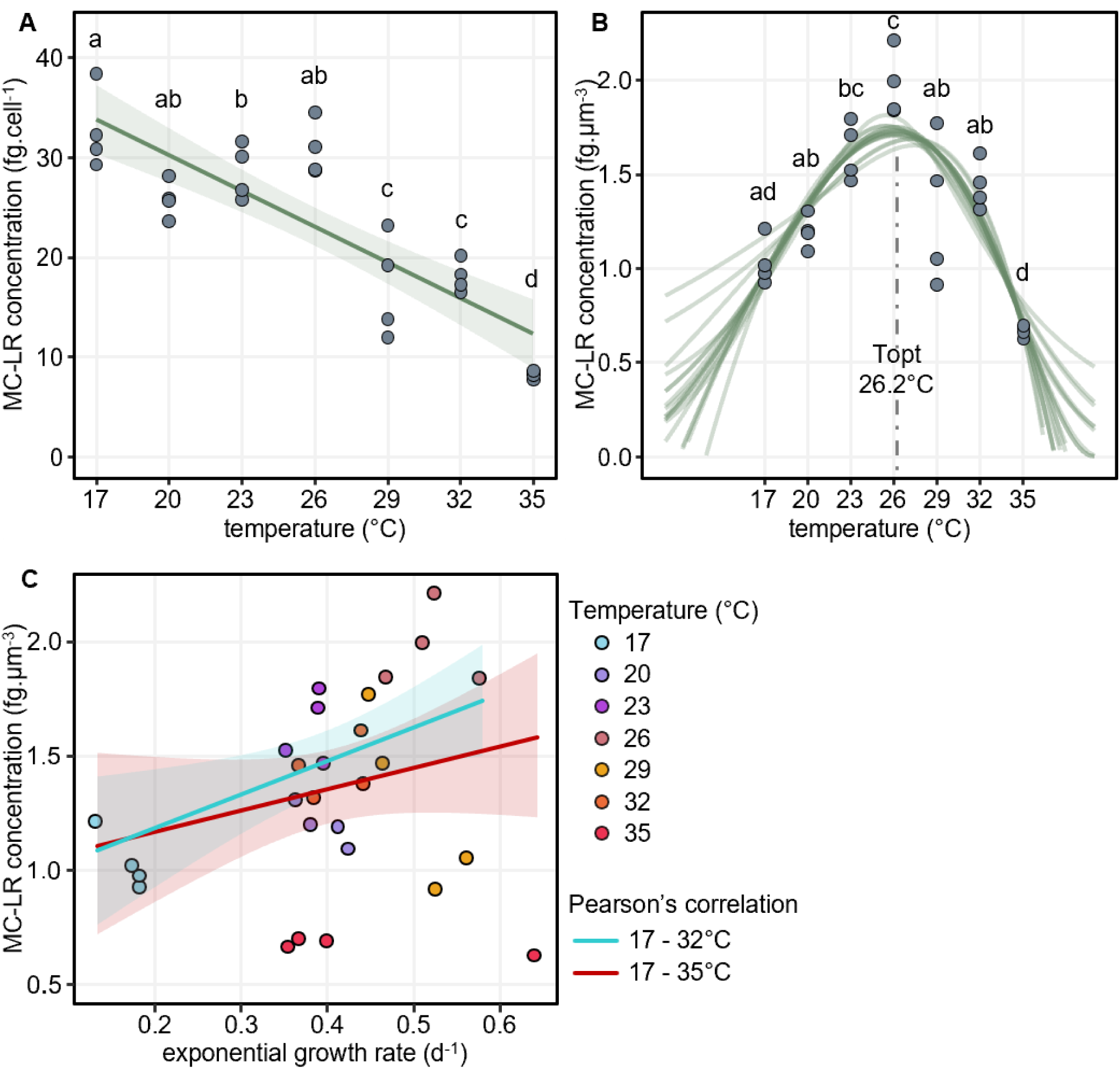
Intracellular microcystin-LR in M. aeruginosa PCC7806 grown at seven temperatures. Points represent the mean for each temperature (n=4). Different letters indicate significant differences between temperature conditions. **(a)** MC-LR Concentration per cell (Table S5; ANOVA, p=7.22e-10). The line represents the linear regression of intracellular MC-LR versus temperature (R^2^=0.7, p=3.6e-8). **(b)** MC-LR Concentration per volume unit (Table S6; ANOVA, p= 1.16e-07). The green lines are the 19 TPC models while the vertical grey dashed line is the mean of optimal temperatures estimated by these models. **(c)** MC-LR concentration per volume unit as a function of the exponential growth rate. Point colors represent the experimental temperature. The blue dashed line indicates the correlation between MC-LR content and growth rate within the 17°C to 32°C range (Pearson’s correlation, p = 0.012), while the red dashed line represents the correlation across the full temperature range, including 35°C (Pearson’s correlation, p = 0.166).

## 4. Discussion

As climate warms and tend to increase the prevalence of cyanobacterial blooms, cyanotoxins are of growing concern for both human and ecosystem health. Understanding how rising mean temperatures will affect microcystin content in cyanobacterial cells is therefore crucial. To address this, we employed an original approach that combined temperature acclimation with cell volume measurements in the assessment of MC-LR content in *Microcystis Aeruginosa* PCC 7806. This work highlights, for the first time, the critical importance of accounting for cell volume when evaluating microcystin content, as we showed that it transformed the observed trend with temperature from a linear decrease to a bell-shaped curve, with the highest value occurring near the thermal growth optimum. These findings provide novel insights into the relationship between temperature and MC-LR content.

When studying the literature, it is difficult to establish the precise relationship between temperature and microcystin content as each study uses different experimental conditions (i.e. different strains of *Microcystis,* temperature, ranging from 17 to 38°C). However, a comprehensive analysis of all data from this literature (since 2015, Figure 4) reveals a general trend of decreasing microcystin content per cell with increasing temperature. Nevertheless, substantial variation in microcystin concentration values and the slopes of these relationships is observed, likely due to methodological differences across studies (Table S7). For instance, at low temperatures the substantial discrepancies observed between our results and those of Peng et al. (2018) could be due to the strain of *M. aeruginosa* used (respectively *M. aeruginosa* PCC7806 in this work and NIES-843 in Peng et al.). It has already been demonstrated that significant variations in microcystin content can occur between different strains within the *Microcystis* genus (Mowe et al., 2015). In addition to differences in the strain used, variations in slope of microcystin content versus temperature observed across studies (Table S8) may reflect interactions between temperature and other environmental factors that differ between studies, such as light intensity (Wiedner et al., 2003) or nutrient availability (Peng et al., 2018). In our case, the lower slope could be attributed to the lower light conditions used in our study (25 µmol photons·m⁻²·s⁻¹ in this study compared to 30-130 µmol photons·m⁻²·s⁻¹ in others), as higher light intensities may amplify the effects of temperature on microcystin content.

**Figure 4.**
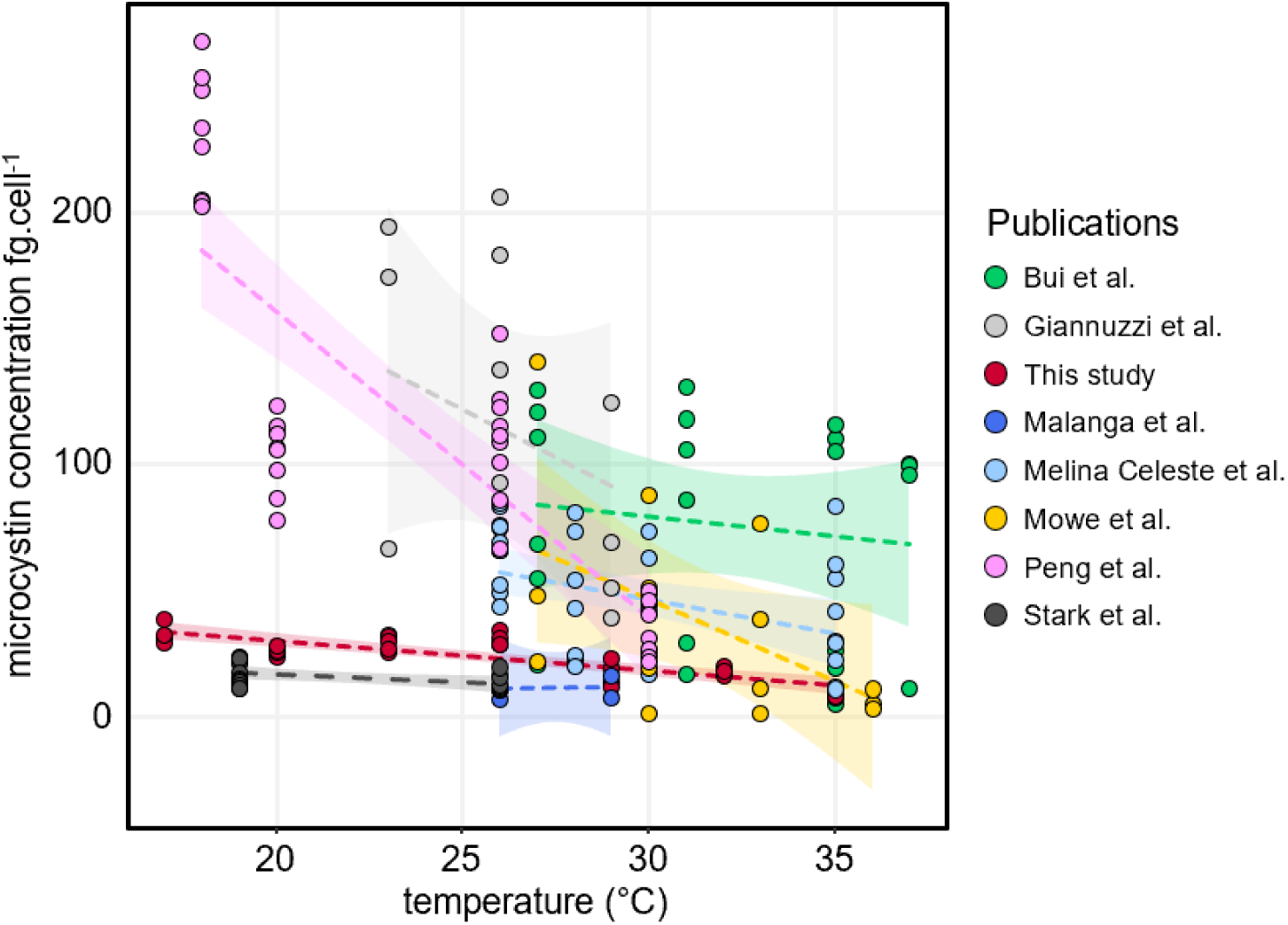
Overview of data from the literature since 2015 on the relationship between intracellular microcystin content and temperature. The dashed lines indicate the trends of cellular microcystin content as a function of temperature for each publication. The color of the points and lines represents the source publication from which the data were extracted: green: Bui et al., 2018; light grey: Giannuzzi et al., 2016; red: Lalloué et al., this study; bright blue: Malanga et al., 2019; light blue: Melina Celeste et al., 2017; yellow: Mowe et al., 2015; pink: Peng et al., 2018; dark grey: Stark et al., 2023. Data were extracted from publications published after 2015 using the online PlotDigitizer application. Only studies investigating the effects of temperature on microcystin content, expressed as fg cell⁻¹, in the Microcystis genus were considered.

However, this trend is based on microcystin content per cell unit, while our results clearly show that *Microcystis* cell volume decreases significantly with increasing temperature, a pattern previously reported in several studies and consistent with indications of a generalized response (Martin et al., 2020; Zohary et al., 2021). Nevertheless, to our knowledge, no study to date has combined microcystin content with cell volume measurements and acclimation to explore the response to increased mean temperature. Microcystin content within a cell is influenced not only by its production and utilization but also by inheritance from the mother cell. This inheritance implies that microcystin content per cell is indirectly affected by temperature. At higher temperatures, cells divide more rapidly (i.e. exhibit higher growth rates) and at a smaller size, as predicted by the temperature-size rule (Angilletta Jr. and Dunham, 2003). Consequently, cells have less time to produce and accumulate microcystin before the next division. However, normalizing by cell volume eliminates this indirect effect, allowing microcystin concentrations to be compared on a standardized basis (i.e. per unit volume), which remains consistent across temperatures. This enables a more accurate assessment of microcystin production and utilization across different thermal conditions. By accounting for cell volume, we found a shift from a linear decrease in MC-LR content with increasing temperature (expressed as fg.cell^-1^) to a bell-shaped distribution (expressed as fg.µm^-3^). The only study which showed a similar trend was that by Rapala and Sivonen (1998) for MC-LR content expressed in mg·g⁻¹, but in *Anabaena spp.* rather than the *Microcystis* genus. These concomitant results on two genera of cyanobacteria, both expressing MC-LR content in biomass (rather than as cell content) in temperature-acclimated cultures, offer a new perspective on the influence of temperature on microcystin content. They allow us to propose two complementary hypotheses to explain the observed trend in microcystin levels which could be attributed either to: 1) differences in its production level, or 2) variations in its utilization (i.e. binding to proteins), depending on the temperature.

The relationship between MC-LR content and temperature follows a classical thermal performance curve shape, suggesting that its levels are governed by the temperature dependence of the enzymatic activity involved in its synthesis, supporting our initial hypothesis. As temperature rises toward the optimum, enzymatic activity is expected to increase, a phenomenon commonly explained by the Arrhenius principle (Schulte et al., 2011). Beyond this optimum, thermal denaturation of enzymes likely leads to a rapid decline in MC-LR production and, consequently, its content. Under our culture conditions, the optimum growth rate was estimated at 27.5°C, in alignment with growth optima reported for the *Microcystis* genus in other studies (e.g., You et al., 2018, 27.5°C). It is noteworthy that, in our study, the estimated optimum temperature for MC-LR content (26.2°C) is close to the optimum for growth. Moreover, while no correlation was observed between growth rate and volumetric MC-LR content across the entire temperature range, a positive correlation emerged between 17°C and 32°C (Figure 3c?). This result suggests a potential decoupling at higher temperatures, possibly due to enzyme denaturation, although this hypothesis remains to be tested. Similarly, Rapala and Sivonen (1998) found a correlation between maximum growth rate and peak microcystin content. Since microcystin synthesis requires substantial nitrogen and energy resources (Omidi et al., 2018), its production could potentially increase near the growth optimum, where cellular metabolic efficiency is assumed to be higher, thereby possibly making more energy available for energetically demanding processes. However, this remains a working hypothesis and warrants further investigation. This raises the possibility that other secondary metabolites, including cyanopeptides, could follow a similar trend, though no direct evidence for this exists in the literature.

The second hypothesis, complementing the first, which could strengthen the bell-shaped distribution, is based on the potential role of microcystins in the temperature stress response. Indeed, MC-LR has been shown to bind to proteins in cyanobacterial cells, forming a protein-bound microcystin pool (Zilliges et al., 2011). The remaining, unused microcystin is referred to as the free microcystin pool. In this study, as in most recent works (Melina Celeste et al., 2017; Peng et al., 2018), a methanol extraction method was used, which exclusively quantifies the free fraction of MC-LR (Meissner et al., 2013). Here, the observed lower free MC-LR content at both higher and lower temperatures may reflect increased binding at these extreme temperatures, suggesting its potential utilization and physiological role under stress conditions. Previous work has shown that MC-LR binds to cysteine residues of redox-sensitive proteins under oxidative stress conditions (Zilliges et al., 2011). This binding is hypothesized to help stabilize the conformation of the key protein, enabling cells to cope with reactive oxygen species induced by adverse environmental conditions such as high light intensity or extreme temperatures (Pospíšil, 2016). However, as this process is irreversible, it may significantly affect intracellular levels of free microcystins (Meissner et al., 2013). This hypothesis was previously proposed by Melina Celeste et al. (2017) for high temperatures alone, and later supported by Roy et al. (2023), who also reported increased MC-LR binding at elevated temperatures. While high temperatures are known to induce oxidative stress, low temperatures can also trigger similar conditions (Kholssi et al., 2023). Thus, the significantly lower MC-LR content observed at 17 and 20°C in our results may extend this hypothesis to low temperatures. Currently, no evidence of MC-LR binding has been reported at low temperatures, underscoring the need for further investigation into MC-LR binding dynamics at suboptimal temperatures, alongside assessments of oxidative stress. This context strengthens the idea that microcystins may play a role in temperature stress response. Indeed, it has been shown that, following a rapid temperature drop, *Microcystis aeruginosa* mutants unable to produce microcystins (PCC7806 ΔmcyB) rely on alternative strategies to protect against reactive oxygen species (Stark et al., 2023). It should be noted that Martin et al. (2020) demonstrated that a rapid temperature drop (from 26°C to 19°C) led to an increase in volume-specific microcystin levels, a result that contrasts with our findings and challenges both of our hypothesis. However, unlike our study, their experiment subjected *Microcystis* cells to an abrupt temperature shift without prior acclimation. In a fluctuating environment, cells must adjust their phenotypes (i.e. acclimate) in response to temperature changes to better cope with new conditions. Since phenotypic plasticity occurs gradually (Fey et al., 2021), it can substantially influence experimental outcomes. This gradual adjustment may complicate the interpretation of microcystin dynamics, as acclimation has already been shown to affect its production (Layden et al., 2022). Moreover, microcystin itself may play a role in the acclimation process (Stark et al., 2023). In this context, the timescale of temperature fluctuations emerges as a critical factor shaping microcystin dynamics and warrants further investigation, particularly as climate change not only raises mean temperatures but also amplifies thermal variability (Kotz et al., 2021). By applying the thermal performance curve framework (to acclimated cultures), this study provides a clearer perspective on how future warming may shape toxic risk in freshwater environments.

Our results demonstrate that temperature significantly influences free microcystin content in *Microcystis aeruginosa* PCC 7806 and, consequently, cell toxicity. With an optimum production/content of free MC-LR observed at a high but still ecologically relevant temperature for temperate lakes (26.2°C) (Maberly et al., 2020), microcystins may become an increasing concern as global temperatures rise. This concern is reinforced by our findings, which suggest that optimal free MC-LR content aligns with the thermal optimum for growth, indicating that the highest toxin levels are likely to occur under conditions that also promote high cyanobacterial biomass. Furthermore, as the optimal temperatures for growth and MC-LR production vary depending on specific culture conditions and Microcystis strains (as discussed earlier), this may broaden the temperature range within which high toxin-related risks can emerge. However, in natural environments, variations in microcystin levels are more complex, as temperature influences also the proportion of toxic versus non-toxic strains within the population. This latter aspect remains a subject of debate within the scientific community, with evidence supporting both an increase (Dziallas and Grossart, 2011) and a decrease (Caen et al., 2024; Ninio et al., 2020) in the prevalence of toxic strains at higher temperatures.

## 5. Conclusions

This study highlights the influence of temperature on free microcystin content in *Microcystis aeruginosa* PCC 7806 and underscores the critical importance of normalizing by its cell volume, as the latter varies with temperature. Here, accounting for cell volume transforms the observed trend from a linear decrease in MC-LR content with increasing temperature into a bell-shaped curve. This pattern may reflect both an increase in production near the optimal temperature for growth and a rise in binding as temperatures approach extremes, inducing stress conditions. By acclimating cells to different temperatures, our results provide a strong basis for understanding how the thermal environment influences microcystin content in the context of global warming. However, as this study focuses on a single strain, further research involving multiple strains, both in monoculture and co-culture, is required to validate and generalize these findings. Additionally, the future toxicity of cyanobacteria warrants further investigation, as climate change not only increases average temperatures but also amplifies thermal fluctuations, which may have additional impacts on microcystin dynamics (Kotz et al., 2021).

## Acknowledgement

This research was funded by the Agence Nationale de la Recherche (ANR), grant ASAP (ANR-22-CE02-0005). We thank Célia Kerbrat for her assistance with microcystin extractions. We also thank Fran Van Wyk de Vries for helping improve the English.

